# Bat things come in threes: within-host dynamics of herpesvirus triple infection in bats

**DOI:** 10.1101/2025.02.05.636680

**Authors:** Samantha Aguillon, Magali Turpin, Gildas Le Minter, Camille Lebarbenchon, Axel O.G. Hoarau, Patrick Mavingui, Muriel Dietrich

## Abstract

Understanding viral community ecology in bats is essential for elucidating shedding patterns and the drivers of co-infections. In this study, we explore the genetic diversity and within-host dynamics of herpesviruses (HSV) in *Mormopterus francoismoutoui*, a tropical insectivorous bat endemic to Reunion Island. Over three consecutive years, we collected saliva samples from seven roosts, including repeated samples from recaptured individuals. Illumina sequencing of HSV PCR-positive samples revealed a high diversity of strains (n = 20), belonging to alpha, beta and gamma-HSV subfamilies. Co-infection was frequent, with 44% of bats shedding strains from all three subfamilies. While most shedding patterns with different subfamilies appeared random, our results suggested a negative influence of gamma-HSV occurrence of the probability of co-shedding alpha-HSV. We also demonstrated a lower HSV diversity in juveniles as compared to adult bats, while pregnancy appeared to increase viral diversity—although this requires further confirmation. Longitudinal recaptures of bats revealed an accumulation of multiple HSV latent-infections over life, as the probability to be infected with a new subfamily increased with time interval between recaptures. Within-host strain dynamics were highly variable, with 79% of bats showing fluctuations in strain diversity over time—either gaining or losing strains—consistent with latency and reactivation mechanisms. These findings provide new insights into the ecological and evolutionary dynamics of herpesviruses in wild bat populations.

## 1. Introduction

Multiple infections with distinct virus populations within a single host are common in wildlife (*e*.*g*. Hoarau et al., 2020) and can occur either simultaneously (co-infection) or sequentially (super-infection) (Alizon, 2013). Simultaneous viral infections can lead to within-host interactions that might be beneficial for viruses, by facilitating genetic exchanges, providing polymerase proteins and enhancing host immune suppression (Shartouny et al., 2022). Conversely, interactions can be antagonistic, by interfering with replication or transmission, via resource competition or interferon-mediated immunity (Delima et al., 2023; Essaidi-Laziosi et al., 2020). In theory, the intensity of within-host interactions is expected to be stronger between closely related parasites due, for example, to overlap in resource use and similarity in the elicited immune recognition profiles (Alizon, 2013; Choisy & de Roode, 2010). Bats constitute a good model to study co-infections as they are hosts of diverse zoonotic viruses and large proportions of co-infected individuals have already been reported in natural populations (*e*.*g*. 42%; Jones et al., 2023; Wang et al., 2023). By altering key epidemiologic factors such as host susceptibility and infection duration (Vaumourin et al., 2015), co-infections could have important consequences for zoonotic spillovers and disease emergence. Understanding how bat viruses may form interactive communities has thus both significant evolutionary and epidemiological implications (Jones et al., 2023).

Herpesviruses (HSV) are a valuable model for studying within-host viral interactions due to their high prevalence in diverse host species and their capacity to establish infection shaped by latency phases and reactivation mechanisms (Cohen, 2020). Herpesvirus family includes three subfamilies with different properties. In humans, alpha-herpesviruses (alpha-HSV) are known for their rapid lytic replication and latency in non-divided cells (neurons), while beta- and gamma-herpesviruses (beta-HSV and gamma-HSV) establish latency in divided cells and have slower replication cycle, which can lead to persistent or chronic infections, especially in individuals with compromised immune systems (Weidner-Glunde et al., 2020). Moreover, the lytic cycle is the principal mechanism for alpha- and beta-HSV, while for gamma-HSV, it is the latency that predominates (Weidner-Glunde et al., 2020). The particularity of herpesvirus to establish chronic infection, combine with important viral population sizes and cycles of reactivation, can lead to significant genetic diversification (Lauring, 2020).

In bats, frequent co-infections have already been reported, with multiple HSV strains and between different subfamilies (beta-HSV and gamma-HSV) (Griffiths et al., 2020; Harima et al., 2023; James et al., 2020; Moreira Marrero et al., 2021). In contrary, infection with alpha-HSV was only described in a few bat species and only reported as single infection (Inagaki et al., 2020; Razafindratsimandresy et al., 2009; Sasaki et al., 2014). Moreover, recent work suggests that maternally-derived antibody protection in juveniles wanes rapidly and that higher prevalence in male adult bats may be shaped by sex-specific behaviour or physiology (Aguillon et al., 2024). However, these studies were focused on prevalence data and only limited research on viral diversity has investigated how host factors could shape multi-infection patterns and within-host dynamics (Griffiths et al., 2022, 2023; Sjodin et al., 2019). In vampire bats (*Desmondus rotundus*) infected with multiple beta-HSV strains, it is suggested that non-competitive strains and latent infections coexist at the population level (Griffiths et al., 2022).

In this study, we investigate the genetic diversity and within-host dynamics of HSV in a tropical insectivorous bat, *Mormopterus francoismoutoui*, endemic to Reunion Island. A recent spatio-temporal study revealed that Reunion free-tailed bats are highly infected by HSV (prevalence of 87%, n = 3,981 bats) with probable latency mechanisms, explaining long-term viral shedding in saliva (Aguillon et al., 2024). We first tested the influence of individual factors (age, sex and reproductive status) on viral diversity, and potential interactions between the co-infecting HSV subfamilies. Within-host dynamics was then assessed, through the recapture of bats, by estimating the probability of changing shedding status (at the subfamily level), and measuring intra-subfamily diversification of HSV strains over time.

## 2. Materials and methods

### 2.1. Field sampling

Longitudinal monitoring of *M. francoismoutoui* in Reunion Island has been ongoing since 2018 (Aguillon et al., 2023a). Saliva samples used in this study were collected in 2018, 2019 and 2020 in 7 roosts and included data from recaptured bats (totalizing between 4 and 5 capture events over the 3 years investigated here).

Briefly, bat capture was performed during the dusk emergence as fully described in Aguillon et al. (2023a), by mainly using harp traps (Faunatech Ausbat) and Japanese mist nets (Ecotone). For each individual, we determined the sex visually and reproductively active status was recorded when females were pregnant and lactating, and in males, when they had large testes (see Aguillon et al., 2023a for more details). In females, we recorded the development of nipples as M0 for non-visible nipples, M1 for visible nipples and M2 for inflated nipples (lactating). Bats were classified as adults or juveniles, by examining the epiphysis fusion in finger articulations. Juveniles were identified when articulations were unfused, a characteristic clearly visible up to seven months of age in this bat species. Beyond this age, some older juveniles may have been mistakenly classified as adults, particularly females without developed nipples (M0). A sterile swab (Puritan Medical Products, USA) was carefully introduced in the corner of the lips to sample saliva and then placed in 250 μL of MEM (Minimum Essential Medium Eagle). Samples were stored in a cool box in the field before to be transferred the same night at -80°C at the laboratory. Finally, we tattooed bats on the right propatagium with an individual alphanumeric code before releasing them in the capture site. Handling of bats was performed using personal protective equipment and gloves were disinfected between each individual bat and changed regularly, and all the equipment was disinfected between sites as well (see protocol in Aguillon et al., 2023a).

### 2.2. Molecular analysis and bioinformatics

DNA extraction from saliva samples and amplification of a fragment of herpesvirus DNA polymerase (207 bp product) were mainly performed as part of Aguillon et al. ‘s study (2024). A nested PCR was used to target a broad spectrum of herpesviruses, including alpha, beta and gamma sub-families (VanDevanter et al., 1996). We took a subset of these data (n = 121 PCR-positive samples / 3,981 tested bats) and added new samples (processed with the same protocol), to finally include PCR-positive samples from both sexes, from adults and juveniles, from bats in active or non-active reproductive stage, and from bats that have been recaptured several times. This corresponds to three biological periods: putative mating (from April to May, majority of males sampled), pregnancy (from October to December, majority of females sampled) and juvenile weaning (from January to March, majority of juveniles sampled). To prepare samples for Illumina sequencing, the second PCR of the nested protocol was repeated with Illumina adapters. PCR amplicons were then processed following an Illumina MiSeq 250 bp paired-end sequencing method at Macrogen Europe (The Netherlands, Amsterdam), using the Herculase II Fusion DNA polymerase Nextera XT Index V2 library kit.

Raw sequence reads were filtered, cleaned and trimmed, removing primers and low-quality reads (unexpected length, missing base) using the FROGS 4.0 pipeline (Bernard et al., 2021; Escudié et al., 2018). Clustering of reads into operational taxonomic units (OTUs) was performed using the SWARM algorithm (Mahé et al., 2014) with an aggregation distance of 5%, following by the removing of chimeras and the filtering of low proportion OTUs (frequency below 0.005%). Resulting OTUs were considered as distinct HSV strains and were checked in GenBank using the Basic Local Alignment Search Tool (BLAST, Altschul et al., 1990) to verify their herpesvirus identity. We checked for appropriate sequencing depth per sample by verifying that the percentage of estimated strain diversity covered in each sample ranged between 95% and 100%, using the function *depth*.*cov* from the R package *hilldiv* (Alberdi et al., 2019). To assess the completeness of our sampling, we also created accumulation curves of strain diversity in each roost, with 95% confidence levels based on 1000 bootstraps, using the R package *iNEXT* (Hsieh et al., 2016) (Fig. S1). Strain diversity was measured based on Hill numbers and *q* = 1, which considers both richness and evenness of taxa.

### 2.3. Phylogenetic analysis

Taxonomic affiliations of HSV-strains into the three subfamilies (alpha-, beta-, gamma-HSV) was performed using a Bayesian tree in BEAST v.2.6.4 (Bouckaert et al., 2019). The tree was built using a Yule model and a HKY site model with invariant and gamma distribution (Hasegawa et al., 1985), after model selection using BIC criterion with JModelTest v2.1.10 (Darriba et al., 2012). We used a reference dataset including sequences previously identified as bat-borne alpha-, beta- and gamma-HSV, retrieved from GenBank, ensuring to select a wide range of host bat families and a diversity of strains within each HSV subfamily. Sequence alignment was constructed using ClustalW (Thompson et al., 2003) and MUSCLE (Edgar, 2004), and then visually checked in CLC sequence Viewer 7.6.1 (Qiagen Aarhus A/S, Aarhus, Denmark). We used a strict molecular clock with a 100 million chain length and sampling every 10^3^ steps, and a burning of 10%. We ran three analyses and combined log outputs (removing 10% of burning for each output) using LogCombiner v2.6.4 (Rambaut & Drummond, 2015). Traces of Markov Chain Monte Carlo (MCMC) were checked for convergence of the posterior using Tracer v1.7.1 (Rambaut et al., 2018). We combined tree outputs (removing 10% of burning for each output) to obtain a consensus tree using LogCombiner v2.6.4 (Rambaut & Drummond, 2015) and then TreeAnnotator v2.6.4 (Rambaut & Drummond, 2019), and finally visualized the consensus tree in FigTree v1.4.4. (Rambaut, 2018).

### 2.4. Statistical analyses

To assess the effect of age (juveniles *vs*. adults) and sex on HSV genetic diversity, we performed generalized linear mixed models (GLMM) including age and sex (and their interaction) as fixed effects, and the bat’s ID as a random effect to account for the recapture of some bats. HSV diversity was measured at the strain level, using both the number of strains (modelled as a Poisson distribution, model M1 in Table S1) and Hill numbers (*q* = 1, modelled as a Gaussian distribution with log link, model M2 in Table S1). The *anova* function with chisquare tests were used to test the statistical significance of explanatory variables (and their interactions) by sequentially removing them from the full models. In addition, to assess strain composition differences between both age classes, we performed a permutational multivariate analysis of variance (PERMANOVA) with 10^3^ permutations, and visualized results with non-metric multidimensional scaling (NMDS) plot using Bray-Curtis dissimilarity index in FROGS 4.1 (Bernard et al., 2021; Escudié et al., 2018).

In order to evaluate the effect of reproductive status (active *vs*. non-active) on HSV genetic diversity (Hill numbers, *q* = 1), we used a gaussian GLMM (with log link) on a subset of data including adult bats during the mating and pregnancy periods. Reproductive status and sex (and their interaction) were included in the model as fixed effects, and the bat’s ID as a random effect (model M3 in Table S1). We re-ran the model after excluding non-pregnant females without developed nipples (M0) to avoid including potential misclassified female juveniles in this analysis (model M3bis). Finally, to investigate potential interactions between HSV subfamilies, we used binomial GLMMs on adult bats, modelling the presence/absence of each subfamily using a binomial distribution (see models M4, M5, and M6 in Table S1). Models included reproductive status and sex (and their interaction), as well as the two other subfamilies as fixed effects, and the bat’s ID as a random effect (with the exception of model M5 where bat’s ID has been removed because of model failure to converge, and thus a GLM was used instead).

We investigated within-host dynamics of shedding on a subset of bats (n = 11) that have been recaptured (n = 45 samples). Specifically, we estimated the probability of acquiring or losing HSV subfamilies through time, by fitting a multinomial logistic regression using the *multinom* function in the *nnet* package, using the R script from Streicker et al. (2019). Conversion status was used as a nominal dependent variable with four levels (no changes, gain, loss, gain and loss) and we defined the non-changing level as baseline outcome against which to compare other shedding status changes (model M7 in Table S1). Finally, we analysed the evolution of HSV genetic diversity at the individual level, by plotting the intra-subfamily strain diversity (as Hill number q=1) over time for each recaptured bat.

All models were constructed and analysed in Rstudio 1.4.1106 (RStudio, 2020) using packages *dplyr, effects, hilldiv, iNEXT, lme4, ggeffects, ggplot2* and *nnet*.

## 3. Results and Discussion

### 3.1. Diversity and abundance of HSV-strains and subfamilies

Based on the analysis of 121 saliva samples, we identified a high diversity of herpesviruses in *M. francoismoutoui*, including 20 strains belonging to the three subfamilies (Fig. 1a). A significant proportion of bats were shedding two (41%) and three (44%) subfamilies. Additionally, 50% of individuals carried between 5 and 8 strains, with some reaching a maximum of 15 strains (Fig. 1b). These findings confirm that HSV co-infections are common in bats (Jones et al., 2023). To our knowledge, this study provides the first report of an alpha-HSV in an insectivorous bat (Gonzalez & Banerjee, 2022) and documents the occurrence of triple HSV infection within a single bat species.

**Figure 1.**
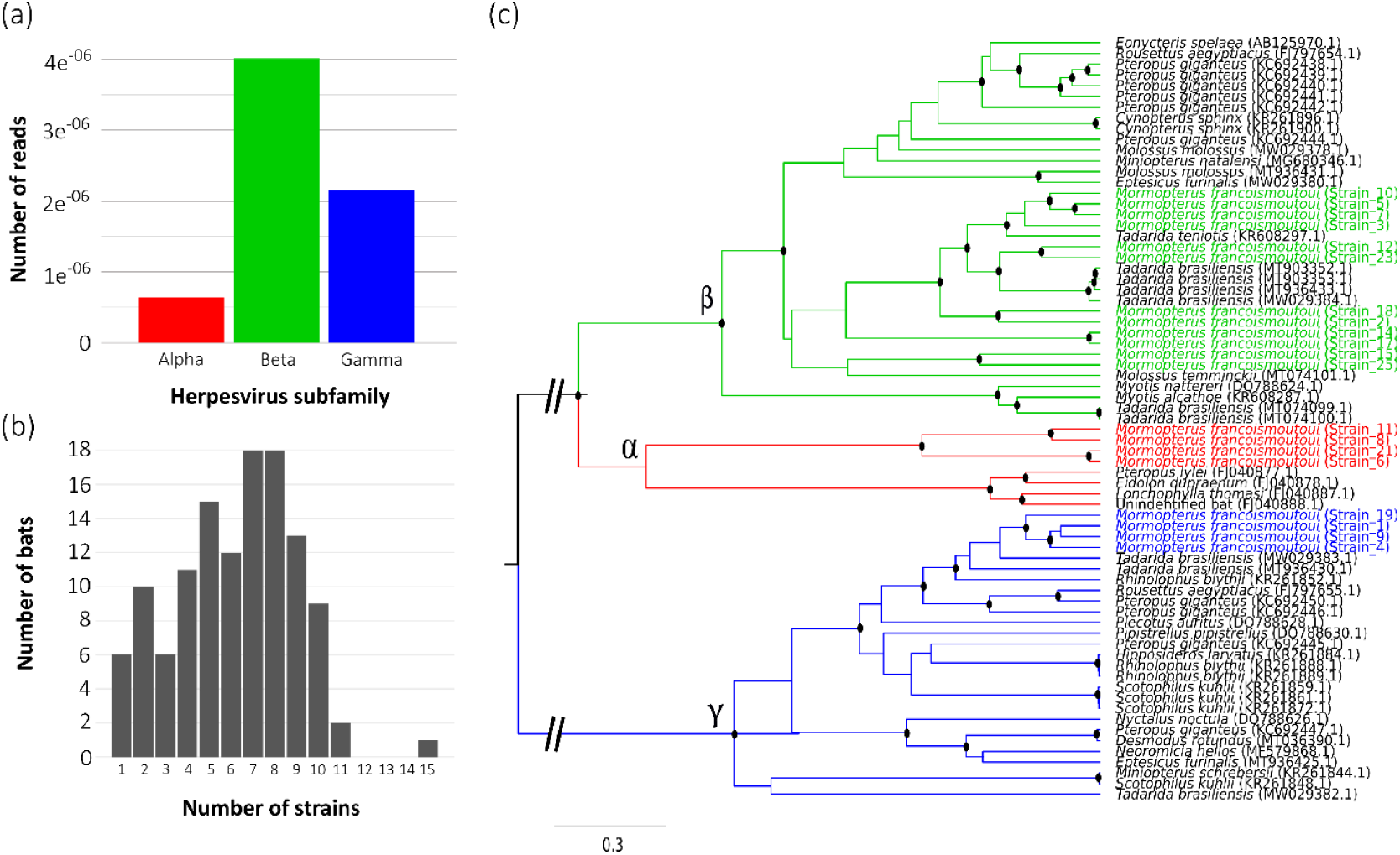
Reunion free-tailed bats are co-infected with herpesviruses from three subfamilies. (a) Abundance of the three herpesvirus subfamilies, based on the total number of Illumina reads over samples. (b) Number of herpesvirus strains hosted by individual bats. (c) Phylogeny shows relationships among herpesvirus strains hosted by *M. francoismoutoui* (n = 20, colored according to the subfamily) and other bat species (n = 63, in black). After alignment, the analysed fragment of the HSV DNA polymerase was 182 bp long. The analysis was conducted using BEAST with the Yule model and the HKY site model, incorporating invariant sites and a gamma distribution. Nodes with posterior probabilities greater than 0.75 are marked with black dots.

Beta-HSV was the most abundant (59% of total reads, Fig. 1a) and diversified (12 strains, Fig. S2) subfamily in *M. francoismoutoui*. Our beta-HSV sequences clustered in several groups closely related to viruses previously found in several molossid bat species, including *Tadarida brasiliensis, Tadarida teniotis* and *Molossus temminckii* (Fig. 1c; Moreira Marrero et al., 2021; Pozo et al., 2016). Gamma-HSV sequences (32% of total reads, Fig. 1a) grouped in a single cluster closely related to HSV from the mollosid bat species *T. brasiliensis* (Moreira Marrero et al., 2021). Alpha-HSV sequences, which were the less abundant (9% of total reads, Fig. 1a), also grouped in a single cluster, and were different from those previously found in different nectar-feeding and fruit bat species (no alpha-HSV reference sequences were available for comparison). The close genetic relationships with HSV from others members of the Mollosidae family (at least for beta and gamma subfamilies) illustrates the probable evolution of host specificity in bat HSVs (Azab et al., 2018; Escalera-Zamudio et al., 2016). Low abundance of alpha-HSV could be explained by the distinct speed in lytic cycles and mechanisms of reactivation among subfamilies, which may reduce the probability of detecting alpha-HSV in saliva, due to their rapid lytic replication (Weidner-Glunde et al., 2020). This is coherent with a modelling study suggesting that active phases are much longer than latency phases for beta-HSV dynamics (Griffiths et al., 2023), that could explain the higher abundance and diversity of beta-HSV observed in *M. francoismoutoui*.

### 3.2. Host-associated determinants of HSV strain and subfamily diversity

Strain diversity was strongly dependent of age, when both measuring the number of strains (model M1: χ^2^_1_ = 20.221, *p* = 7^e-06^, Table S1) (Fig. 2a) and Hill numbers (model M2: χ^2^_1_ = 14.118, *p* = 2^e-04^, Table S1) (Fig. S3a). The PERMANOVA test also detected a significant difference in strain composition between adults and juveniles (F = 0.03, *p* < 0.01), with reduced diversity observed in juveniles (Fig. 2b). Indeed, the majority of juveniles (8 over 9) were shedding only one HSV subfamily (beta- or gamma-HSV, Fig. S3b) and a reduced mean number of strains (predicted mean = 2.51, Fig. 2a). Only one juvenile was shedding the three subfamilies (Fig. S3b). In comparison, almost half of adults were shedding the three subfamilies simultaneously (52 over 112, Fig. S3b) and a higher number of strains (predicted mean = 6.44, Fig. 2a). Interestingly, Griffiths et al. (2022) did not find an effect of age on the number of HSV strains in *Desmodus rotundus*. This contrasting result could be due to species-specific difference, or because our juvenile class was restricted to newborns up to 4 months-old bats, while Griffiths et al. (2022) also included subadults. These subadult bats had probably been exposed to a higher HSV diversity, supported by our previous study showing that juveniles get infected rapidly (Aguillon et al., 2024). Thus, in addition to reduced HSV prevalence previously reported in juvenile bats (*e*.*g*. Aguillon et al., 2024; Dietrich et al. 2018; Griffiths et al., 2020), our findings revealed a reduced genetic HSV diversity in young juveniles, as compared to adult bats. This supports the hypothesis that maternal antibodies provide early protection, followed by multiple latent HSV infections accumulating over the bat’s lifespan.

**Figure 2.**
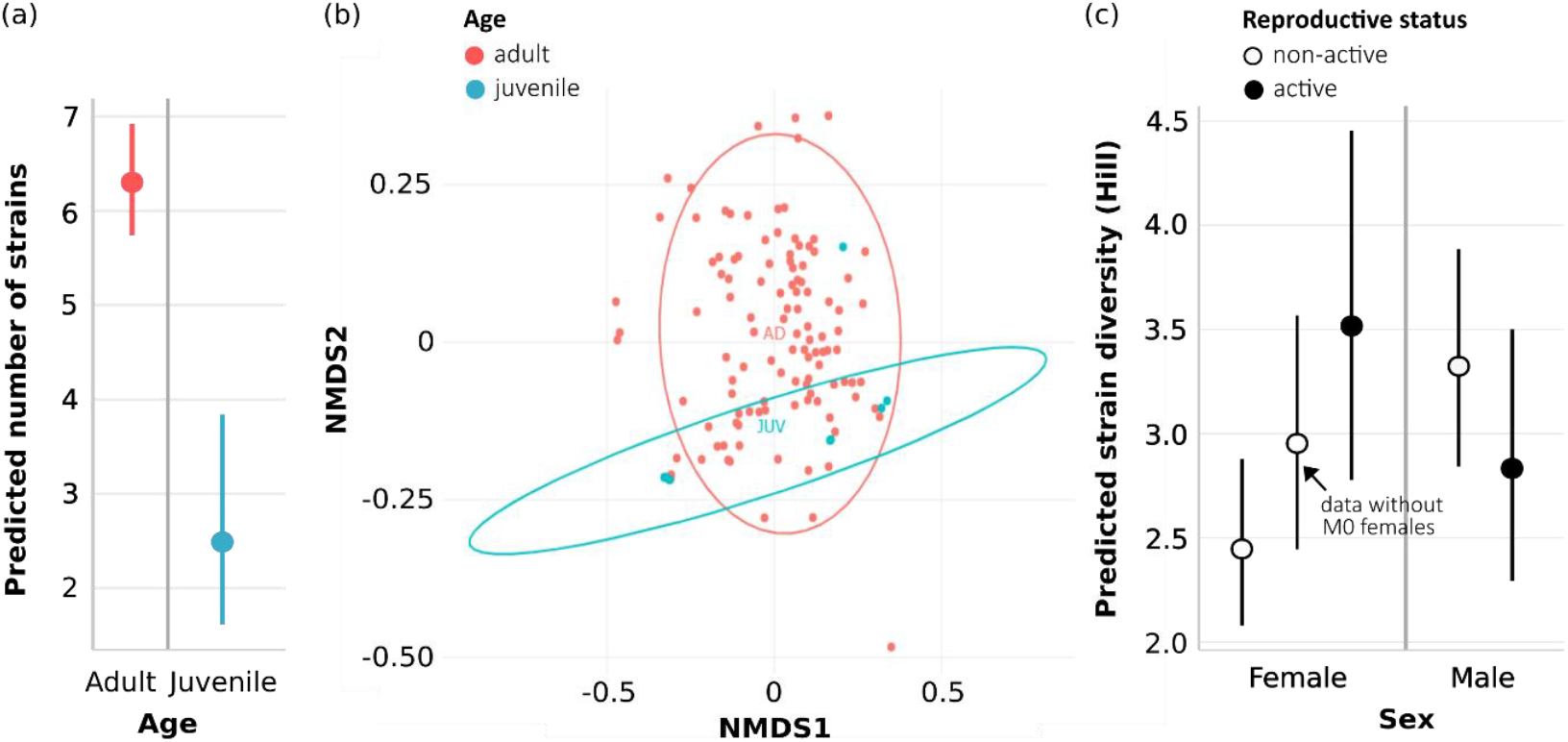
Reduced herpesvirus diversity in juveniles and apparent increased diversity in pregnant females of *M. francoismoutoui*. (a) Predicted number of strains according to bat’s age (model M1, Table S1). (b) NMDS plot of herpesvirus strain composition according to bat’s age. (c) Predicted strain diversity (Hill number, *q* = 1) according to sex and reproductive status of adult bats (model M3, Table S1). For females, strain diversity was also modelled after excluding non-pregnant females without visible nipples (M0) to avoid including potential misclassified female juveniles (model M3bis, Table S1).

In adult bats, we did not detect a global effect of sex on HSV strain diversity (model M3: χ^2^_1_ = 2.279, *p* = 0.131, Table S1). This is consistent with a recent study on *Desmodus rotundus* (Griffiths et al. 2022), although others have shown that sex effect may be variable among bat species (Sjodin et al., 2020). However, we observed a positive effect of the reproductive status, dependent of the sex (model M3: χ^2^_1_ = 9.186, *p* = 0.002, Table S1), as pregnant adult females were shedding a significantly higher HSV diversity compared to the non-pregnant ones (Fig. 2c). A similar pattern has been found in Puerto Rican bats (Sjodin et al., 2020), which suggests that reactivation of herpesvirus shedding during pregnancy leads to a diversification of the within-host viral community. However, when excluding non-pregnant females with non-visible nipples (M0), the predicted strain diversity in non-reproductively active adult females increased, and the interaction between sex and reproductive status was no longer observed (model M3bis: χ^2^_1_ = 3.668, *p* = 0.055, Table S1). This suggests that the pregnancy effect we detected in adult females may be more related to the age of bats, as some non-pregnant M0 females, classified as adults, could in fact be old juveniles (about 11 months-old) that were still shedding a lower HSV diversity than true adults.

Finally, prevalence of beta-HSV was not influenced by the co-occurrence of alpha-HSV (model M5: χ^2^_1_ = 3.389, *p* = 0.066, Table S1) and gamma-HSV (model M5: χ^2^_1_ = 1^e-05^, *p* = 0.997, Table S1). This indicates that patterns of co-shedding between beta-HSV and the two others subfamilies were random, supporting low levels of cross-protective immunity for these pairwise HSV combinations (Griffiths et al., 2022). However, adult bats shedding gamma-HSV had less probability to shed alpha-HSV at the same time (model M4: χ_1_ = 4.137, *p* = 0.042, Table S1) (Fig. 3a). Such negative interaction has already been reported in bats, but only for HSV strains within the beta-subfamily (Anthony et al., 2013). Further investigations on the infection mechanisms of HSV in bats are needed to explore factors leading to these subfamily-specific interaction patterns.

**Figure 3.**
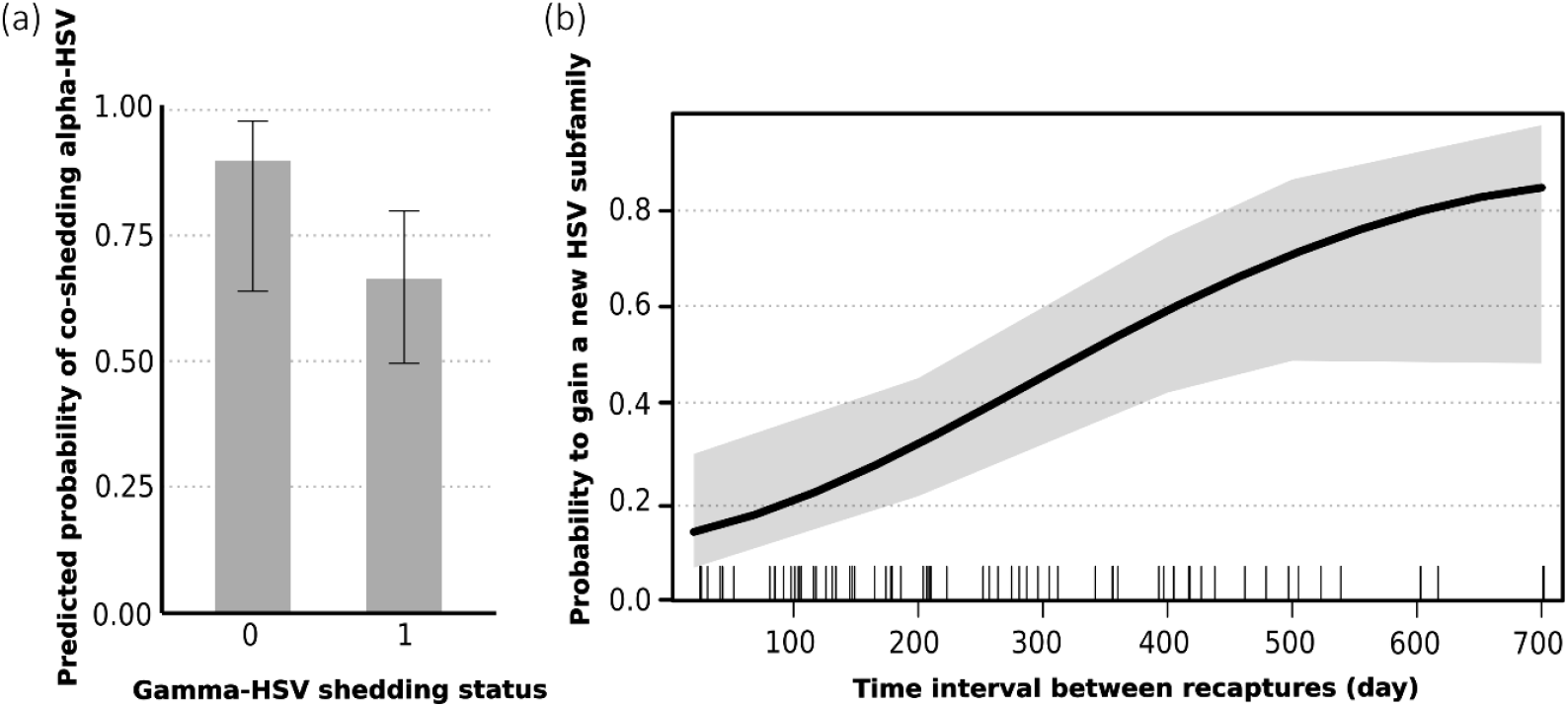
Interaction among herpesvirus subfamilies and their accumulation over life in *M. francoismoutoui*. (a) Predicted negative interaction of gamma-herpesvirus on alpha-herpesvirus co-shedding probability (model 4, Table S1). (b) Probability to gain a new herpesvirus subfamily over time (model 7, Table S1). Dashes across the x-axis indicate recapture of bats and the shading area represent the 95% CI on the mean probability (solid black line).

### 3.3. Within-host dynamics of HSV

Among 11 adult males recaptured after 25 to 702 days, only two were initially HSV-negative (including one being juvenile at the first capture) and all were then systematically shedding over recapture events. For the majority of individuals (52.86%), bats were shedding the same HSV subfamily over recapture events, although the gain of one or two subfamilies was also largely observed (38.57%). Therefore, we reported few loss or gain/loss events (8.57%). The different categories of transition in HSV shedding status had a different probability to occur over time (model M7: χ^2^_3_ = 16.06, *P* < 0.001). Indeed, the probability that bats acquire a new HSV subfamily increased over time (Fig. 3b), while the probability to loss or gain/loss was low and stable over time (Fig. S4). These results support that HSV establish latent infection in bats, as suggested by previous results on prevalence data with bats remaining positive through time (Aguillon et al., 2024; Griffiths et al., 2023).

At the HSV subfamily level, investigation of within-host evolution of genetic diversity revealed highly dynamic strain acquisition/release processes (Fig S5). Indeed, 78.57% of bats both gained and loose strains between captures, and 14.29% of bats only gained strains over time. These frequent shifts are consistent with long-term HSV persistence and may be the result of latency phases and lytic reactivation (Griffiths et al. 2022), although it does not exclude the occurrence of clearance processes as well.

## 4. Conclusion

Our work illustrates that herpesviruses provide an original study system to analyse within-host viral community dynamics in bats. The widespread occurrence of triple HSV infections (alpha, beta and gamma) in *M. francoismoutoui* shed light on individual factors influencing within-host viral diversity and interactions between viruses. While the precise mechanisms underlying HSV persistence and reactivation remain complex and only partially understood, further research is needed to assess how anthropogenic stressors influence HSV shedding dynamics. Moreover, elucidating the mechanisms of HSV reactivation and diversification in bats is particularly important, as these infections may facilitate the co-shedding of other potentially zoonotic pathogens, such as paramyxoviruses and *Leptospira* (Aguillon et al., 2024). Understanding these processes could provide critical insights into the broader implications of viral ecology and spillover risks at the human-wildlife interface.

## Ethics

Bat capture and manipulation techniques were evaluated by the ethic committee of Reunion Island, approved by the Ministère de l’Enseignement Supérieur, de la Recherche et de l’Innovation (APAFIS#10140-2017030119531267), and conducted under a permit (DEAL/SEB/UBIO/2018-09) delivered by the Direction de l’Environnement, de l’Aménagement et du Logement (DEAL) of Reunion Island.

## Data accessibility

Sequences have been deposited in NCBI under the SRA accession number PV034612 to PV034631 and metadata are available in Zenodo (10.5281/zenodo.14810050). Supplementary material is available online.

## Authors’ contributions

S.A.: Conceptualization, Formal analysis, Investigation, Visualization, Writing - original draft. M.T.: Formal analysis, Methodology, Writing - review & editing. G.LM.: Investigation, Writing - review & editing. C.L.: Investigation, Writing - review & editing. A.H.: Investigation, Writing - review & editing. P.M.: Supervision, Writing - review & editing. M.: Conceptualization, Formal analysis, Funding acquisition, Investigation, Methodology, Supervision, Validation, Writing - original draft.

## Acknowledgements

We are grateful to Eco-Med Océan Indien, Biotope, the DEER of Région Réunion (Direction de l’Exploitation et de l’Entretien des Routes), the DRT of Département Réunion (Direction des Routes et des Transports). We are grateful to Clara Castex, Timothé Chenin, Jérémy Dubrulle, Avril Duchet, Léa Joffrin, Riana Ramanantsalama, Pablo Tortosa, Céline Toty, Guillaume Verchère for their assistance in the field.

