## Supplementary materials for "Bat things come in threes: within-host dynamics of herpesvirus triple infection in bats"

**Table S1. Summary of the statistical models used to analyse herpesvirus genetic diversity in *M. francoismoutoui*.** Models include all tested variables, with interaction indicated by an asterisk, and significant variables in bold. The percentage of deviance explained by the final model was calculated after comparison to a null model. M0: female with no visible nipples. All the models include Bat's ID as a random effect (except model M5).

| *Type and model number* | *Analysed samples* | *Response variable* | *Distribution* | *Explanatory variables* | *Xi²* | *p* | *Deviance explained (%)* |
|---|---|---|---|---|---|---|---|
| GLMM M1 | All individuals<br>n = 121 | Number of strains | Poisson | Age * Sex<br>Sex<br>**Age** | 0.038<br>1.949<br>**20.221** | 0.846<br>0.163<br>**7e-06** | 3.48 |
| GLMM M2 | All individuals<br>n = 121 | Hill diversity (strain level) | Gaussian (log) | Age * Sex<br>Sex<br>**Age** | 0.189<br>1.373<br>**14.118** | 0.664<br>0.241<br>**2e-04** | 4.26 |
| GLMM M3 | Adults bats<br>Pregnancy and mating periods<br>n = 96 | Hill diversity (strain level) | Gaussian (log) | **Repro * Sex**<br>Sex<br>Repro | **9.186**<br>2.279<br>7e-04 | **0.002**<br>0.131<br>0.979 | 4.74 |
| GLMM M3bis | Adults bats<br>Pregnancy and mating periods, without non-pregnant M0<br>n = 82 | Hill diversity (strain level) | Gaussian (log) | Repro * Sex<br>Sex<br>Repro | 3.668<br>2e-04<br>0.150 | 0.055<br>0.990<br>0.698 | |
| GLMM M4 | Adults bats<br>Pregnancy and mating periods<br>n = 96 | Alpha prevalence | Binomial | Repro * Sex<br>**Gamma**<br>Beta<br>Sex<br>Repro | 0.045<br>4.137<br>2.881<br>1.084<br>0.152 | 0.833<br>**0.042**<br>0.090<br>0.298<br>0.696 | 3.88 |
| GLM M5 | Adults bats<br>Pregnancy and mating periods<br>n = 96 | Beta prevalence | Binomial | Repro * Sex<br>Gamma<br>Alpha<br>Sexe<br>Repro | 5e-09<br>1e-05<br>3.389<br>1.663<br>3.712 | 0.999<br>0.997<br>0.066<br>0.197<br>0.054 | |
| GLMM M6 | Adults bats<br>Pregnancy and mating periods<br>n = 96 | Gamma prevalence | Binomial | Repro * Sex<br>Beta<br>Alpha<br>Sexe<br>Repro | 1.832<br>0<br>0.280<br>0.020<br>0.029 | 0.176<br>1<br>0.597<br>0.888<br>0.865 | |
| M7 | Recaptured bats<br>n = 70 (capture events) | Conversion of subfamily | Multinomial | **Time** | 16.06 | 0.001 | 11.78 |

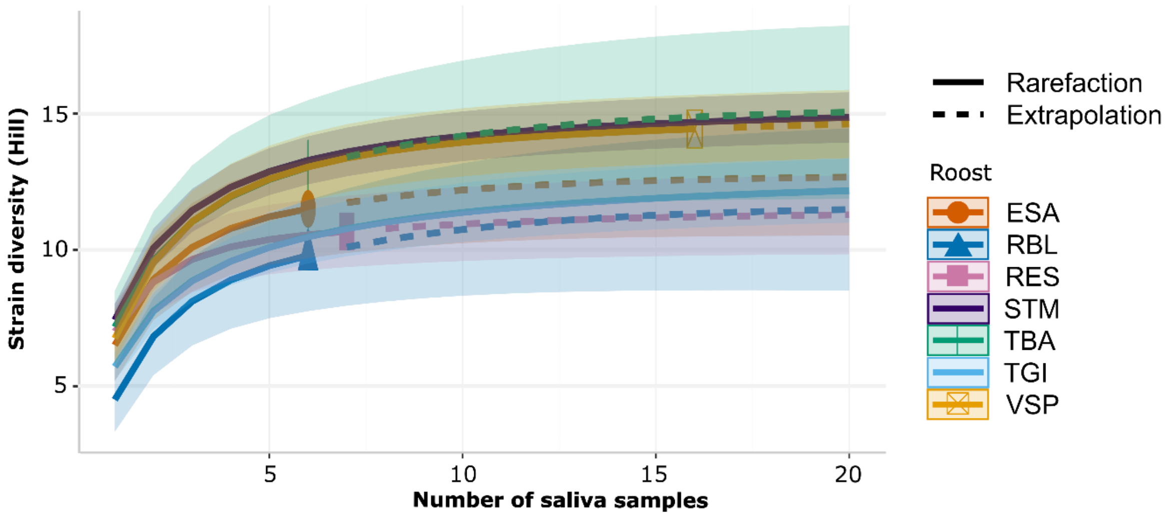


**Figure S1. Accumulation curves of herpesvirus strain diversity (Hill, *q* = 1) according to the number of saliva samples collected per roost.** The curve was constructed based on 1000 bootstraps and the 95% confidence interval is represented by shaded areas.

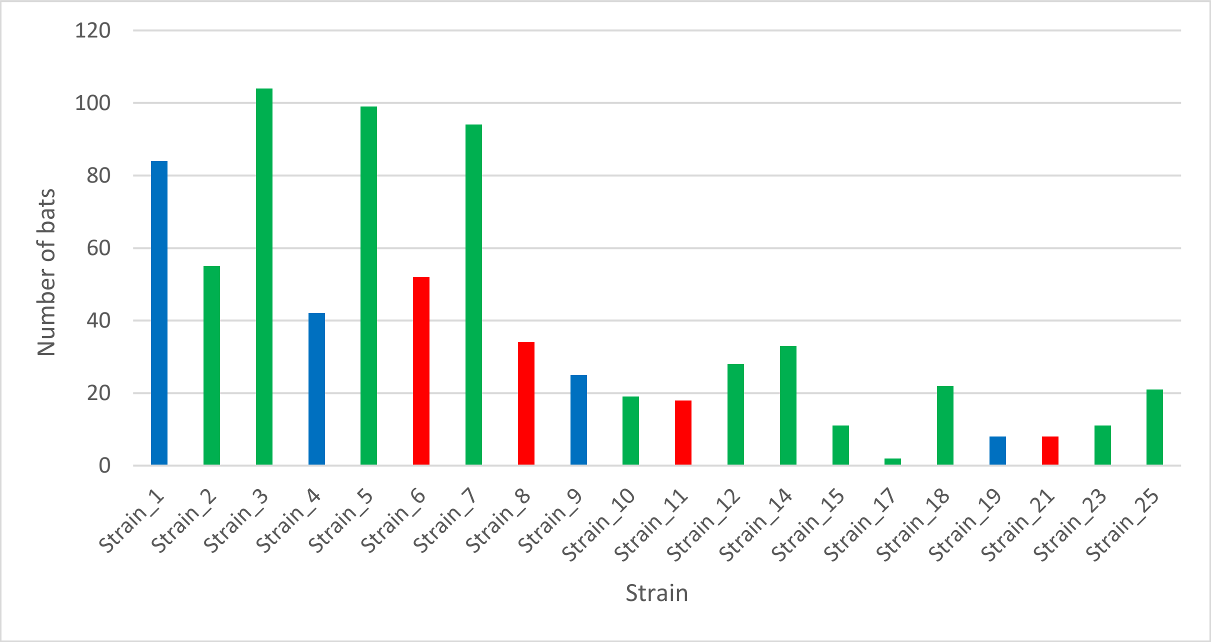


**Figure S2. Abundance of the 20 herpesvirus strains identified in *M. francoismoutoui*.** Colors correspond to the three HSV subfamilies (alpha in red, beta in green, gamma in blue). Abundance is determined by the total number of Illumina reads across all samples.

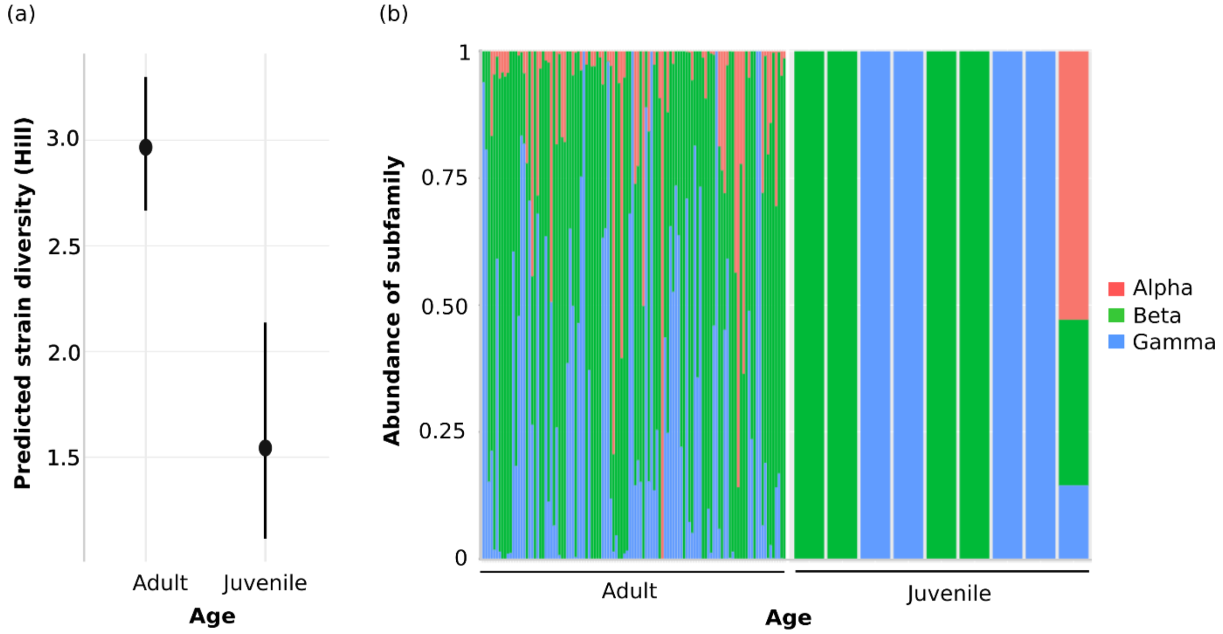


**Figure S3. Diversity of herpesvirus strains and abundance of subfamilies in *M. francoismoutoui* according to bat's age.** (a) Predicted number of strains in adult and juvenile bats according to model M1 (Table S1). (b) Distribution of herpesvirus subfamilies (color-coded) in adult and juvenile bats. Each bat is represented by a vertical column.

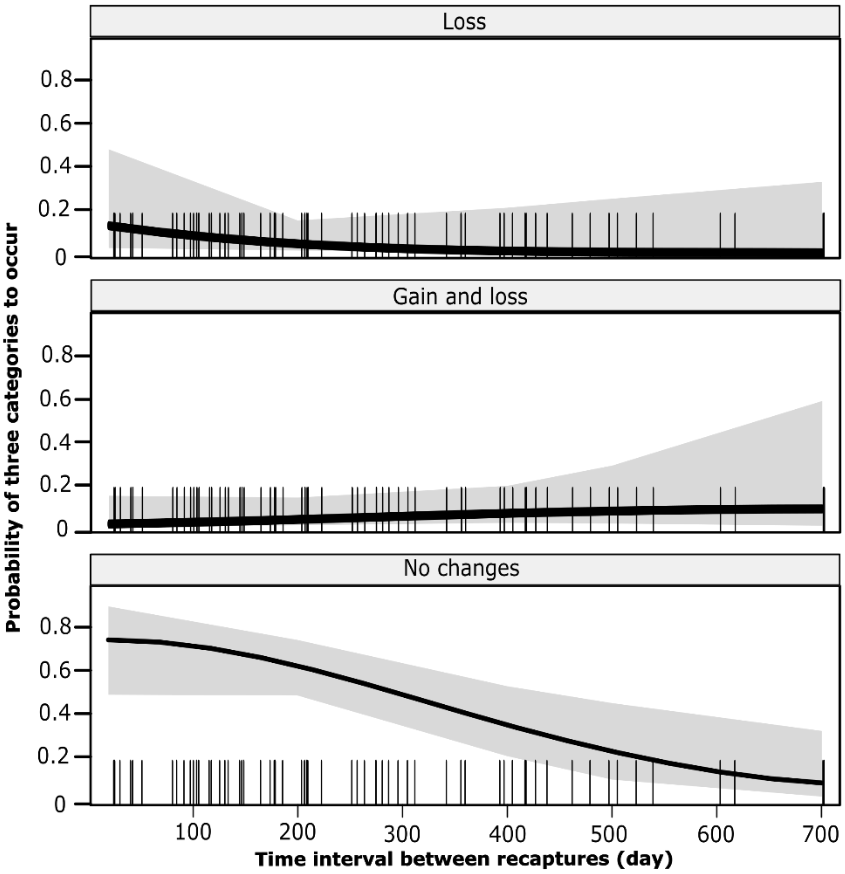


**Figure S4. Probability of the three categories of transition in shedding status**. Values were based from predictions from model M7, with 95% confidence intervals in grey. Probability of the fourth category (gain) is presented in Figure 3b.

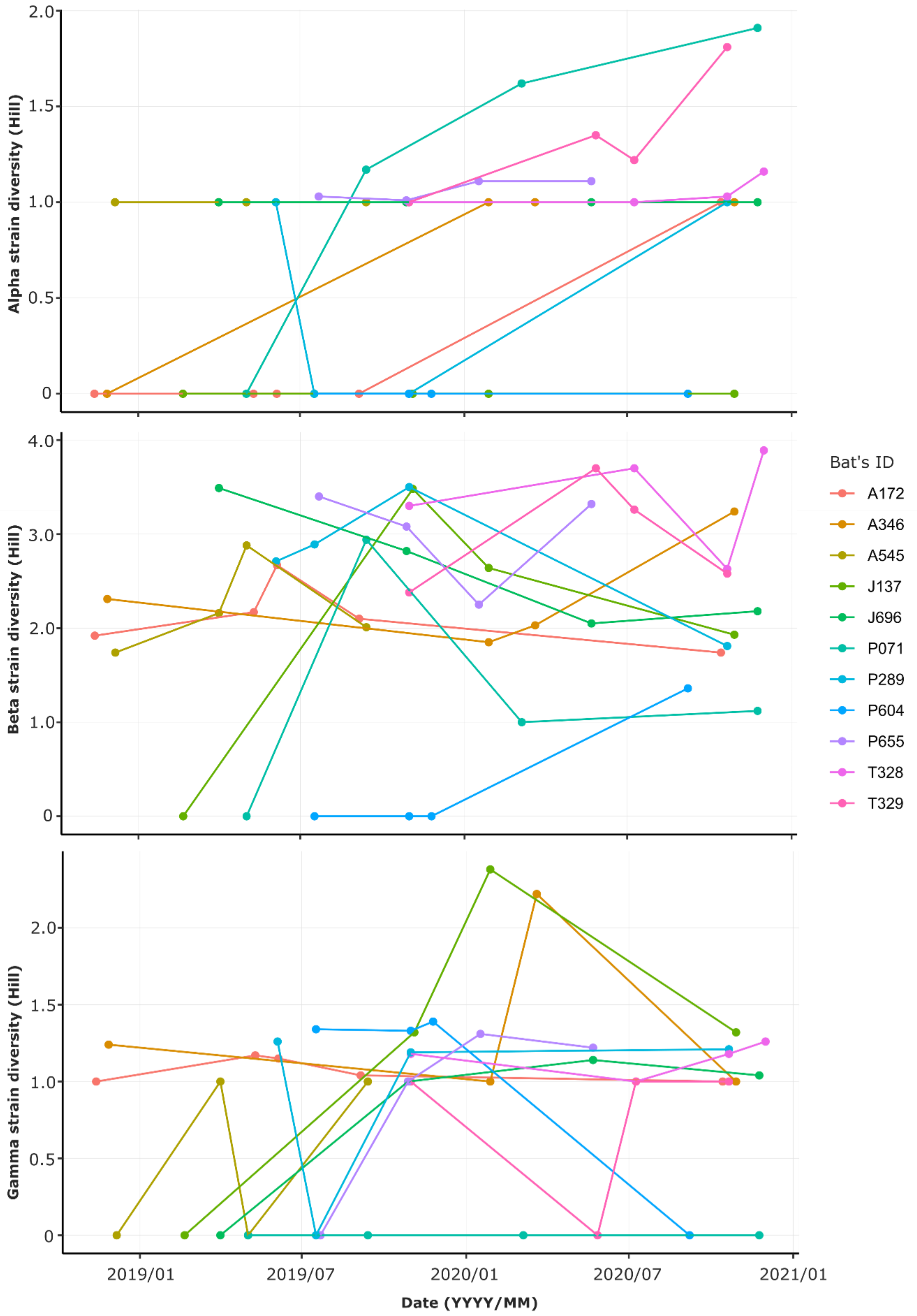


**Figure S5. Temporal evolution of herpesvirus strain diversity at the individual bat level, for alpha-, beta- and gamma-herpesvirus subfamilies.** Strain diversity was calculated based on Hill numbers ($q$ = 1) of herpesvirus strains over time for 11 individual bats captured at least four time.
